# Concurrent administration of COVID-19 and influenza vaccines enhances Spike-specific antibody responses

**DOI:** 10.1101/2023.09.12.557347

**Authors:** Susanna E. Barouch, Taras M. Chicz, Ross Blanc, Domenic R. Barbati, Lily J. Parker, Xin Tong, Ryan P. McNamara

**Affiliations:** Ragon Institute of MGH, MIT, and Harvard

**Author notes:** These authors contributed equally to this work (Susanna E. Barouch, Taras M. Chicz, Ross Blanc, Domenic R. Barbati).

**Keywords:** bivalent, COVID-19, influenza, vaccine, heterologous vaccine, Omicron, SARS-CoV-2, XBB.1.5

## Abstract

The bivalent COVID-19 mRNA boosters became available in fall 2022 and were recommended alongside the seasonal influenza vaccine. However, the immunogenicity of concurrent versus separate administration of these vaccines remains unclear. Here, we analyzed antibody responses in healthcare workers who received the bivalent COVID-19 booster and the influenza vaccine on the same day or different days. IgG1 responses to SARS-CoV-2 Spike were higher at peak immunogenicity and 6 months following concurrent administration compared with separate administration of the COVID-19 and influenza vaccines. These data suggest that concurrent administration of these vaccines may yield higher and more durable SARS-CoV-2 antibody responses.

## INTRODUCTION

The bivalent COVID-19 mRNA vaccines encoded ancestral and BA.5 Spike ^1^, and subsequent Omicron lineages emerged that further escaped antibody recognition ^2^ including XBB strains ^3,4^. The rollout of the bivalent COVID-19 mRNA vaccines in fall 2022 coincided with the seasonal influenza vaccines. However, it has remained unclear how concurrent administration of COVID-19 mRNA and influenza vaccines may impact antibody profiles generated.

Here we profiled antibody responses of healthcare workers who received the bivalent COVID-19 mRNA booster and the seasonal influenza vaccine on the same day or different days. We observed higher IgG1 responses to multiple Spike variants both at peak immunogenicity and 6 months in individuals who received the COVID-19 and influenza vaccines on the same day compared with those who received the vaccines on separate days. Other IgG subclasses and antibody isotypes did not display a similar trend. Our study suggests an immunological benefit to concurrent vaccination with COVID-19 mRNA and seasonal influenza vaccines.

## METHODS

### Experimental Outline and Study Participants

Participants were enrolled as a part of the Massachusetts Consortium on Pathogen Readiness (MassCPR) with informed consent. Individuals were divided into participants who received an influenza vaccine on the same day as the bivalent COVID-19 mRNA vaccine or those who received the two vaccines on different days within 4 weeks. Vaccines were administrated September-December 2022. Serum samples were obtained 3-4 weeks and 6 months after the COVID-19 booster.

### Antibody Profiling

Antibody subclasses, isotypes, and Fc-receptor binding antibodies were assayed for binding to antigens listed in Supplementary Table 1 as described elsewhere^5^. The breadth of antibody subclass and isotype binding was quantified by standardizing each subclass and isotype to Wu-1 Spike binding for receiving the vaccinations on different days (Supplementary Figure 1).

### Quantification and Statistical Analysis

All figures and statistics were done using R Studio V 6.0. For correlation plots, a Spearman’s Rank correlation was calculated against individual pairings and plotted as a heatmap.

## RESULTS

### Concurrent bivalent COVID-19 and influenza vaccination led to higher Spike IgG1 responses

A cohort of 42 healthcare workers was followed longitudinally after bivalent COVID-19 mRNA boosting in fall 2022. Sera were evaluated at weeks 3-4 after boosting (peak immunogenicity) and at month 6 after boosting. The cohort was divided into individuals who received the COVID-19 booster and the influenza vaccine on the same day (n = 12) or different days (n = 30) (**Figure 1A**).

**Figure 1.**
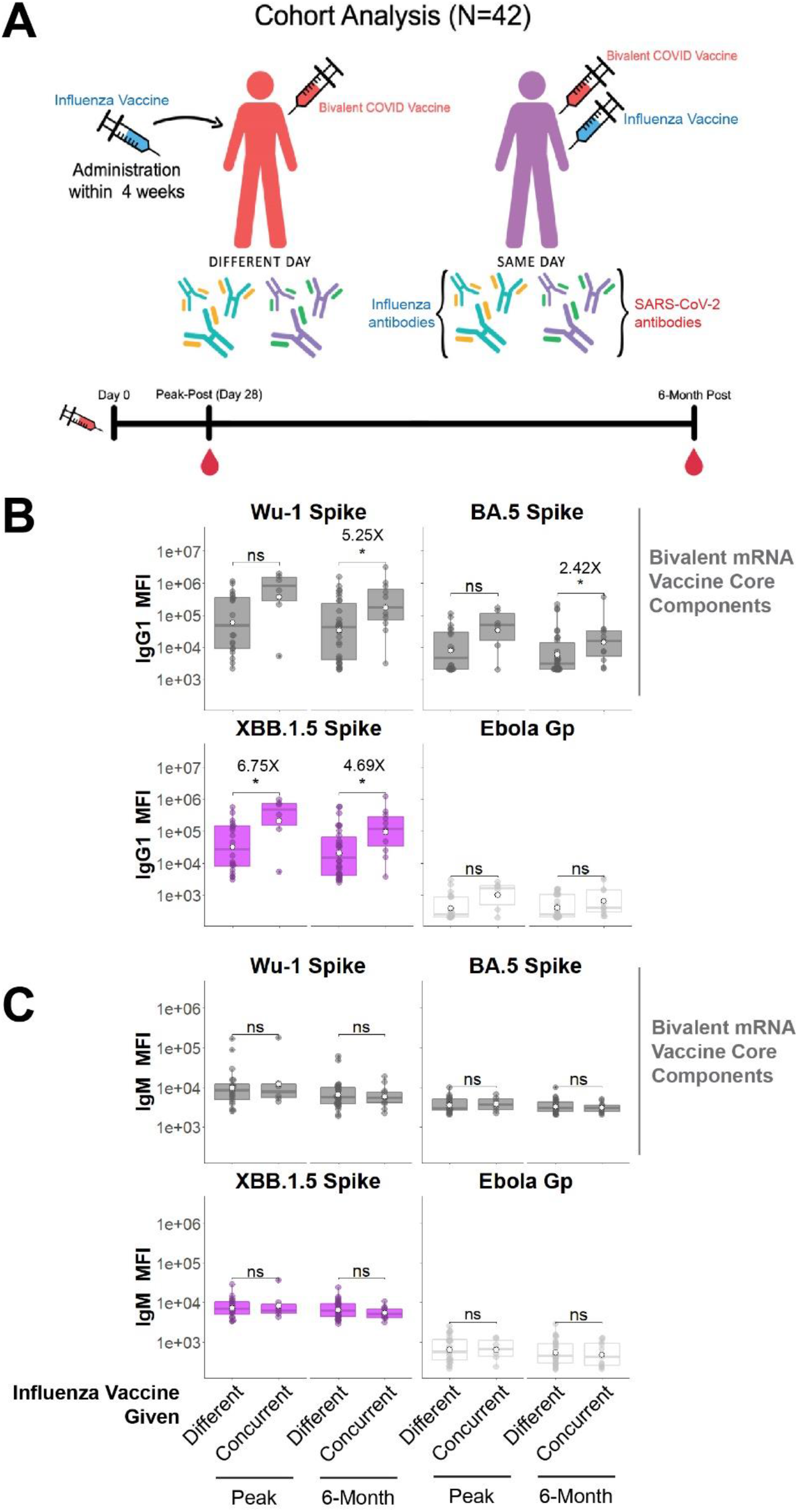
Concurrent bivalent COVID-19 mRNA and influenza boosters induce more durable IgG1 responses to Spike. A. Cohort used in this study. Participants were divided into those who received a bivalent mRNA COVID-19 booster and flu vaccine on the same day (concurrently) or different days. Blood was drawn at peak immunogenicity (2-4 weeks) and 6 months after the bivalent COVID-19 mRNA booster. B. IgG1 antibody responses to the COVID-19 Spike vaccine immunogens ancestral (Wu-1) and Omicron BA.5 (grey), the two components of the bivalent mRNA vaccine, as well as one of the most prevalent SARS-CoV-2 variants of the time XBB.1.5 (purple, primary comparison group). Ebolavirus glycoprotein (Gp) was used as a negative control (white). White circles indicate the means. Fold differences for statistically significant comparisons are shown. For all comparisons, * = p<0.05, ns = not statistically significant, Mann–Whitney U test **/** Wilcoxon rank-sum test. C. IgM antibody responses were quantified similarly to (B) and serves as a control to assess *de novo* antibody affinity maturation.

IgG1 responses to the vaccine immunogens Wu-1 and BA.5 Spike at 6 months were 5.25 and 2.42 fold higher, respectively, in individuals who received the bivalent COVID-19 booster and influenza vaccines concurrently compared with separately (**Figure 1B – gray bars**). IgG1 responses also trended higher at peak immunogenicity at weeks 3-4 with concurrent administration. IgG1 responses to XBB.1.5 Spike was significantly higher at peak (6.75 fold) and at 6 months (4.69 fold) in individuals who received the COVID-19 and influenza vaccines concurrently (**Figure 1B** – **purple bars**). In individuals who received the vaccines on different days, no differences were observed based on vaccination order (data not shown). No IgG1 responses were observed to Ebolavirus glycoprotein (negative control).

No differences were observed in IgM responses (**Figure 1C**), suggesting that concurrent COVID-19 and influenza vaccination drove enhanced recall responses, not *de novo* responses ^5^. IgG2, IgG3, IgG4, and IgA subclasses also showed no differences between groups (Supplementary Figure 2).

### Concurrent bivalent COVID-19 and influenza vaccinations increased IgG1 breadth 6 months post-vaccination

We next assessed if concurrent vaccination increased IgG1 binding breadth to SARS-CoV-2 Spikes. These included Alpha, Beta, Delta, Gamma, BA.1, BA.5, BQ.1.1, and XBB.1.5 Spikes. Comparisons between concurrent and different vaccination days showed consistently increased IgG1 responses and sustained FcγRIIIA responses at 6 months to all these Spike variants in individuals who received the vaccines concurrently (**Figure 2**). We also observed a more robust correlation between IgG1 and IgG3 with FcγRs at both peak immunogenicity and 6 months in individuals who received the vaccines concurrently compared with those who received the vaccines on different days (Supplementary Figure 3).

**Figure 2.**
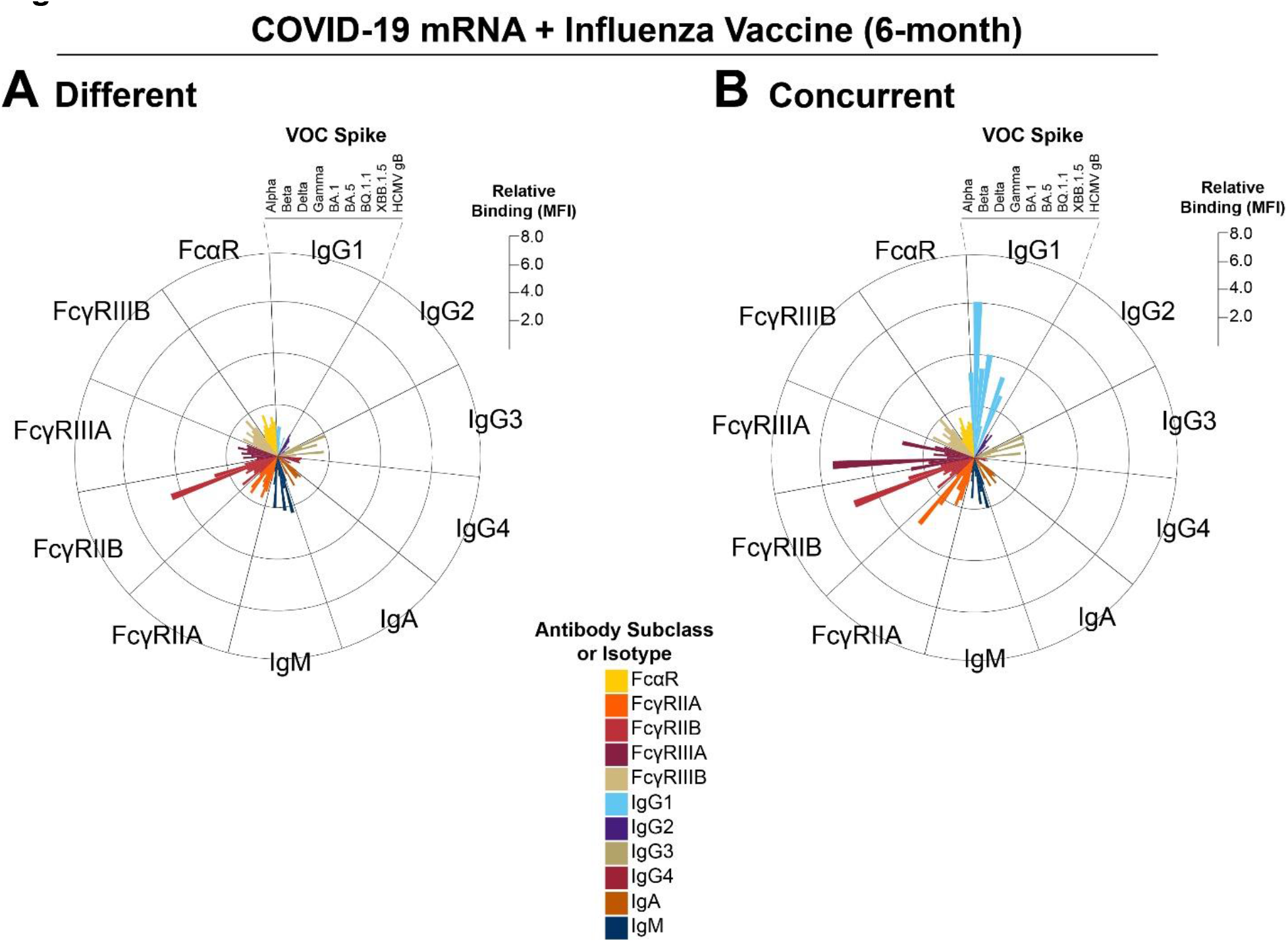
Concurrent COVID-19 mRNA and influenza vaccination selectively expand IgG1 binding breadth to multiple Spike variants. A. Radar plot showing relative binding of individual antibody isotypes, subclasses, Fc-gamma receptor (FcγR), and Fc-alpha receptor (FcαR) binding antibodies to identified SARS-CoV-2 and control antigens. Individual bars represent the median fluorescence activity (MFI) of a specific feature standardized to that antibody subclass/isotype response to Wu-1 Spike for individuals who received the bivalent mRNA and influenza vaccine on different days 6 months after bivalent COVID-19 booster. The scale on the right represents fold MFI increase relative to Wu-1 Spike for each antibody feature. B. Radar plot showing relative binding of individual antibody isotypes, subclasses, FcγR, and FcαR-binding antibodies to the identified SARS-CoV-2 and controls antigens for individuals who received the bivalent COVID-19 and influenza vaccine concurrently. Individual bars represent the MFI of a specific feature standardized to that antibody/isotype response to Wu-1 Spike for individuals who received the bivalent mRNA and influenza vaccine on different days 6 months after the bivalent COVID-19 booster. The use of Wu-1 Spike responses of individuals who received the vaccines on different days was used as a standard to compare across groups. The scale on the right represents fold MFI increase. All antibody isotypes, subclasses, and FcR-binding antibodies are shown in distinct colors, and a legend is shown at the bottom.

No significant differences were observed for antibody responses to nucleocapsid, arguing against infection impacting humoral profiles (Supplementary Figure 4A). In contrast with Spike, no significant differences were observed in antibody responses to influenza hemagglutinin in individuals who received the bivalent COVID-19 vaccine and the influenza vaccine concurrently or separately (Supplementary Figure 4B).

## DISCUSSION

This study shows that concurrent administration of the bivalent COVID-19 booster and the inactivated influenza vaccine on the same day resulted in higher Spike-specific IgG1 antibody responses at peak and 6 months compared with administration of these vaccines on separate days. Safety profiles of concurrent COVID-19 and influenza vaccination have been reported ^6^, but limited data exists on durable antibody responses following different vaccination schedules. One previous report analyzing quadrivalent influenza and mRNA-1273 vaccines showed no antigen interference or safety concerns but did not evaluate detailed long-term antibody responses to these vaccines ^7^.

IgG1 is the most abundant serum IgG subclass and is capable of both neutralizing and non-neutralizing functions. A correlate of protection against COVID-19 of neutralizing antibodies has been reported, but this was only studied for the ancestral Wu-1 virus ^8^. Other reports have suggested that Fc-effector functions may also be required for protection against Omicron variants Spike ^5,9,10^.

One limitation of our study is the relatively small size of this cohort, which primarily consisted of healthcare workers. Therefore, this cohort may not be reflective of the general population given discrepancies in age ranges, sex, and occupational exposure risks. In addition, timing in this study was defined as relative to COVID-19 vaccination. As such, we may not have captured peak influenza responses as accurately as we did for SARS-CoV-2.

In summary, our results suggest potential benefits of concurrent administration of the COVID-19 mRNA vaccines and the seasonal influenza vaccine for induction of Spike-specific IgG1. Because of the expected seasonality of SARS-CoV-2 and influenza, both vaccines will likely continue to be recommended. Our results suggest that concurrent administration of these vaccines should be considered as a strategy to potentiate IgG1 responses to the COVID-19 vaccine and possibly improve vaccine effectiveness ^8,11^.

## AUTHOR CONTRIBUTIONS

Conceptualization: S.E.B., X.T., and R.P.M.

Data curation: S.E.B., T.M.C., D.B., R.B., L.J.P., and R.P.M.

Formal analysis: S.E.B., T.M.C., D.B., R.B., L.J.P., and R.P.M.

Funding acquisition: R.P.M.

Investigation: S.E.B., T.M.C., D.B., R.B., L.J.P., X.T., and R.P.M.

Methodology: S.E.B., T.M.C., D.B., R.B., L.J.P., X.T., and R.P.M.

Project Administration: X.T. and R.P.M.

Resources: R.P.M.

Software: S.E.B., L.J.P., and R.P.M.

Supervision: X.T. and R.P.M.

Validation: S.E.B., T.M.C., D.B., R.B., L.J.P., and R.P.M.

Visualization: S.E.B. and R.P.M.

Writing – original draft: S.E.B, T.M.C., D.B., R.B., and R.P.M.

## ACKNOWLEDGEMENTS

We thank Nancy Zimmerman, Mark and Lisa Schwartz, an anonymous donor (general financial contribution to the Ragon Institute), and Terry and Susan Ragon for their donations to the Ragon Institute. The Systems Serology Laboratory of Ryan P. McNamara is funded for work in this manuscript by the Massachusetts Consortium of Pathogen Readiness (MassCPR) and the National Institutes of Health (1P01AI165072-01, 1P01AI168347-01, 4U01CA260476-02, CIVIC 75N93019C00052) and the Bill and Melinda Gates Foundation (INV-031624 and INV-001650). The conclusions of the results presented here are those of the authors and do not purport to represent those of the funders and funding agencies.

