## Supplementary Information for "Concurrent administration of COVID-19 and influenza vaccines enhances Spike-specific antibody responses"

Supplementary Information includes Figures S1-S4 and their corresponding captions, and Supplementary Table 1.

**Supplementary Figure 1**

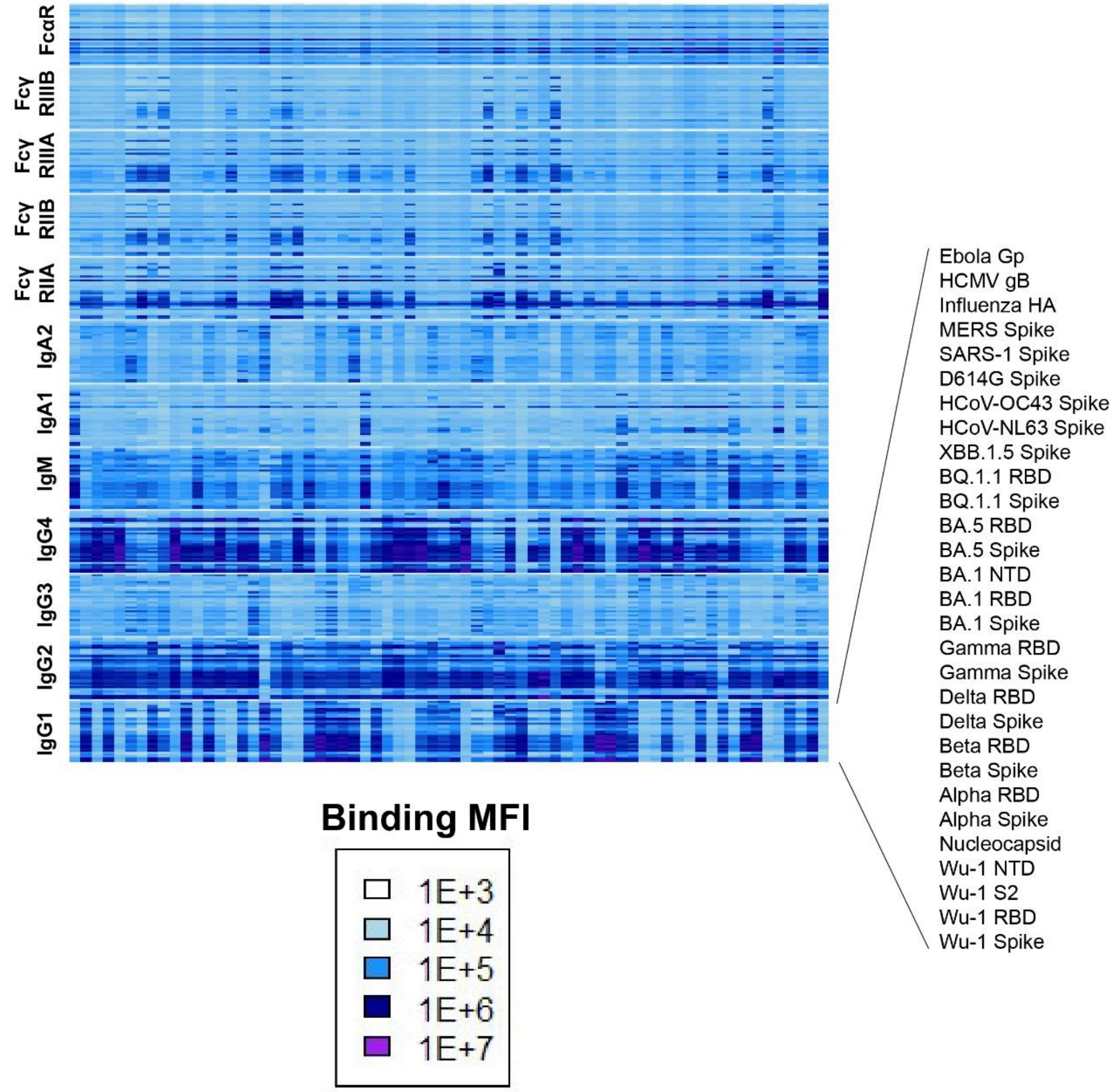


**Supplementary Figure 1. Heatmap of antibody binding to antigens.**
Binding profiling was done for antibody isotypes, subclasses, and FcRs shown on the left. Each column represents a single anonymized individual. Each row represents the antibody's binding to the antigen (order list shown on the right). Binding was quantified through median fluorescence intensity (MFI) as arbitrary units, and a scale is shown at the bottom.

### Supplementary Figure 2

**A**

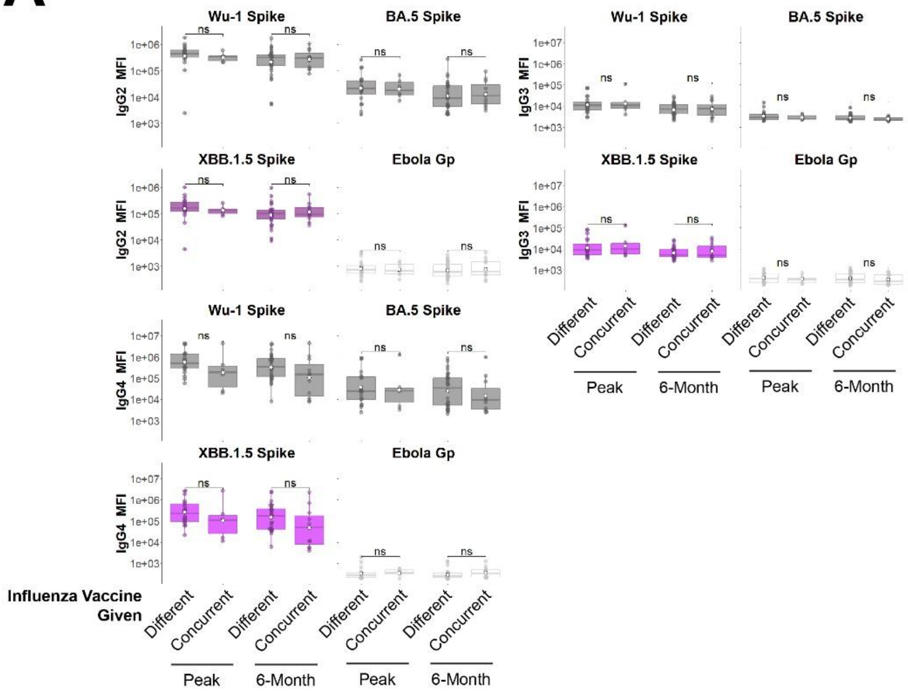


**B**

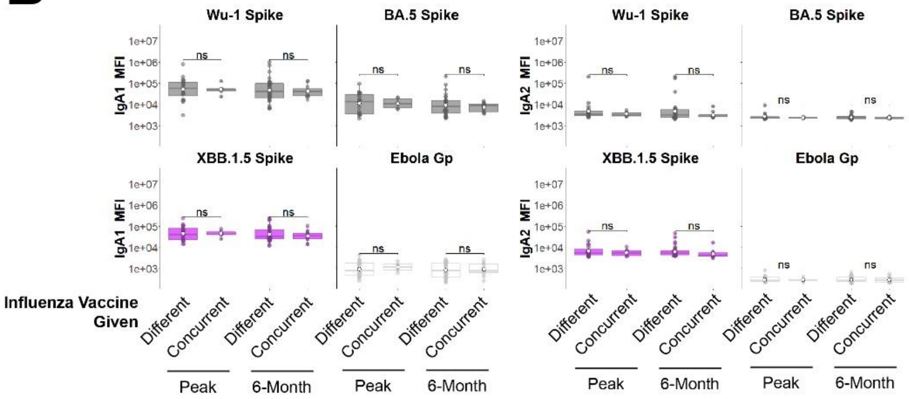


**Supplementary Figure 2. Binding antibodies other than IgG1 display no difference in binding capabilities from different or concurrent COVID-19 and influenza vaccinations.**

A. IgG2, IgG3, and IgG4 binding to ancestral (Wu-1), Omicron BA.5, Omicron XBB.1.5, and Ebola Gp were quantified for the indicated groups at peak immunogenicity and 6 months post-vaccination.

B. Same as A, but for IgA1 and IgA2.

For all comparisons, * = $p<0.05$, ns = not statistically significant, **Mann–Whitney U test / Wilcoxon rank-sum test.**

**Supplementary Figure 3**

A

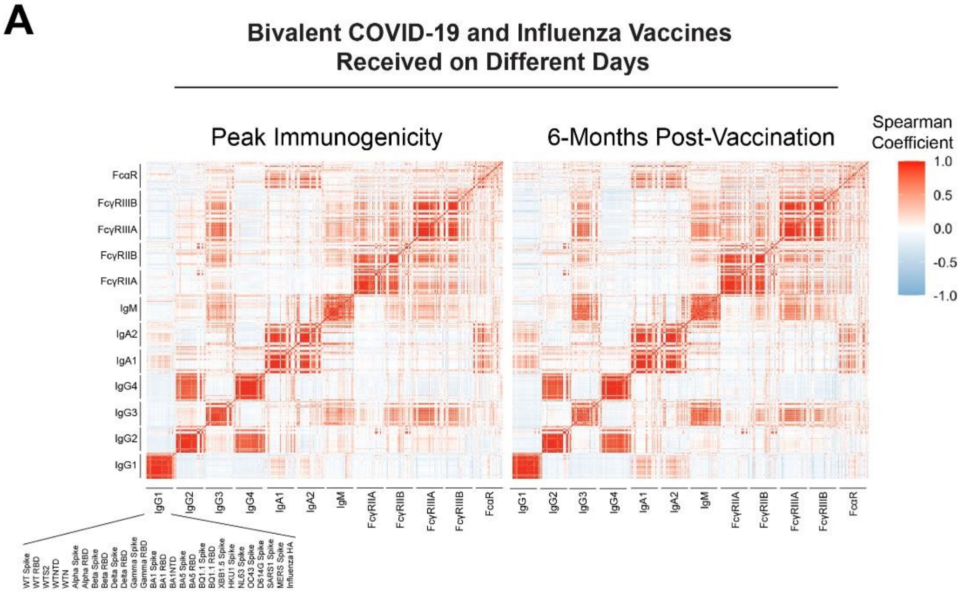


B

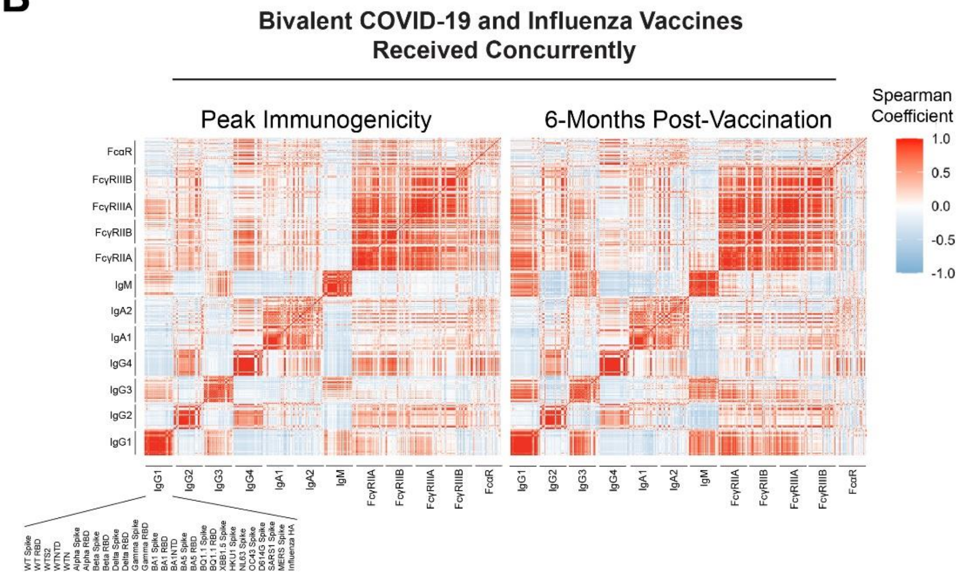


**Supplementary Figure 3. Correlation heatmaps of individuals show higher coordination between Fab-binding and FcR-binding antibodies for individuals who received the bivalent COVID-19 and influenza vaccines concurrently.**

A. Spearman's correlation heatmap of individual antibody binding features with the indicated antigens at peak immunogenicity (left) and 6 months post-vaccination (right) for individuals that received the bivalent COVID-19 and seasonal influenza boosters on different days. Shown on the right is the heatmap legend for the correlation index for each pairwise comparison.

B. Same as (A), but for individuals that received the bivalent COVID-19 and seasonal influenza vaccine concurrently. The Heatmap legend is shown on the right.

### Supplementary Figure 4

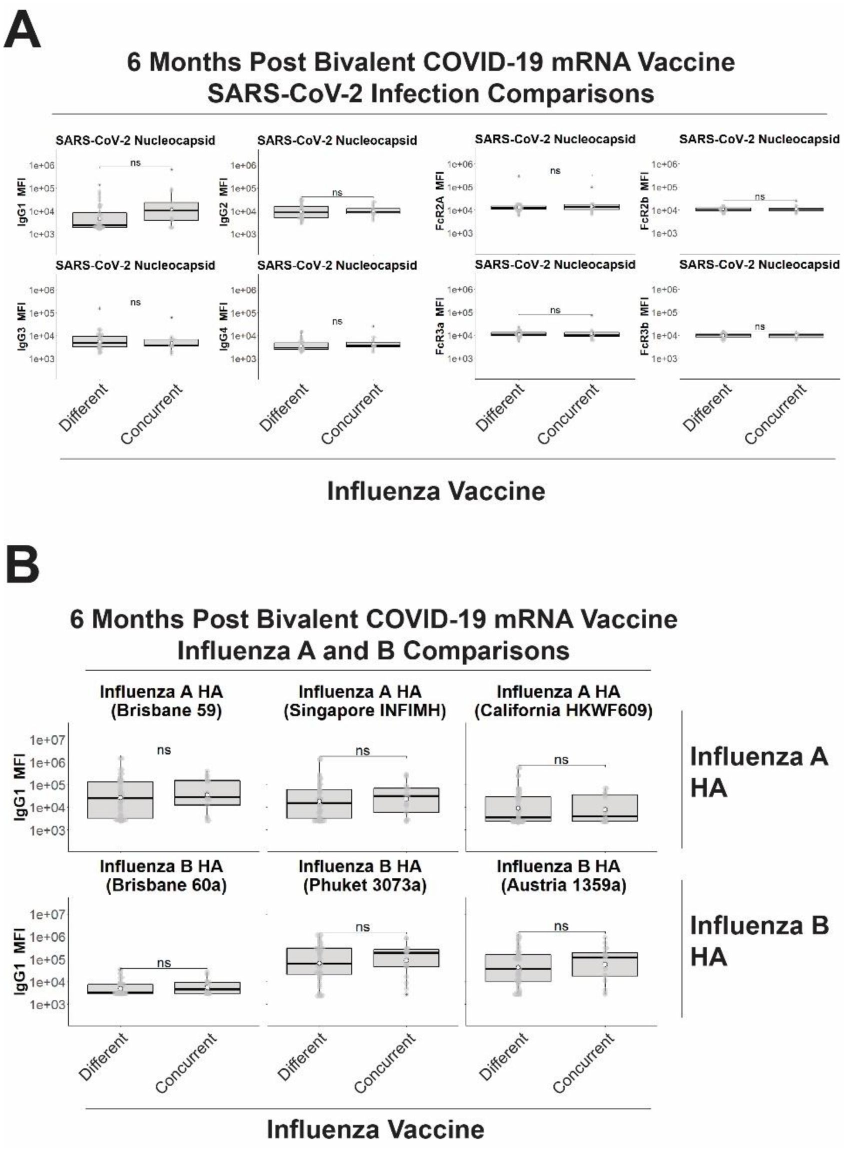


**Supplementary Figure 4. Infections during the observation period are not driving antibody profile distinctions between groups.**

A. IgG1-IgG4 and FcγRIIA – FcγRIIIB binding to nucleocapsid was quantified for individuals who received the bivalent COVID-19 and influenza vaccines on different days or concurrently. Nucleocapsid was used as a bait antigen as it is not a component of the mRNA vaccine. Shown are responses at the 6 month time window to capture any indications of infection throughout the study period.

B. IgG1 responses towards influenza A (top row) and influenza B (bottom row) hemagglutinin antigens (HA) for individuals who received the bivalent COVID-19 and influenza vaccines on different days or concurrently. Shown are responses at the 6 month time window. For all comparisons, * = $p<0.05$, ns = not statistically significant, **Mann–Whitney U test / Wilcoxon rank-sum test.**

### Supplementary Table 1

| REAGENT or RESOURCE | SOURCE | IDENTIFIER |
|---|---|---|
| **Antibodies** | | |
| Mouse anti-human IgG1-PE | SouthernBiotech | 9054-09 |
| Mouse anti-human IgG2 PE | SouthernBiotech | 9070-09 |
| Mouse anti-human IgG3-PE | SouthernBiotech | 9210-09 |
| Mouse anti-human IgG4-PE | SouthernBiotech | 9200-09 |
| Mouse anti-human IgA1 | SouthernBiotech | 9130-09 |
| Mouse anti-human IgM | SouthernBiotech | 9020-09 |
| **Chemicals, peptides, and recombinant proteins** | | |
| SARS-CoV-2 WT Spike | Sino Biological | 40589-V08H4 |
| SARS-CoV-2 WT S1 | Sino Biological | 40591-V08H |
| SARS-CoV-2 WT RBD | Sino Biological | 40592-V08H |
| SARS-CoV-2 WT S2 | Sino Biological | 40590-V08B |
| SARS-CoV-2 WT NTD | Sino Biological | 40591-V49H |
| SARS-CoV-2 D614G Spike | Sino Biological | 40589-V08B4 |
| SARS-CoV-2 Alpha Spike | Sino Biological | 40589-V08H12 |
| SARS-CoV-2 Alpha RBD | Sino Biological | 40592-V08H82 |
| SARS-CoV-2 Beta Spike | Sino Biological | 40589-V08B7 |
| SARS-CoV-2 Beta RBD | Sino Biological | 40592-V08H59 |
| SARS-CoV-2 Gamma Spike | Sino Biological | 40589-V08B10 |
| SARS-CoV-2 Gamma RBD | Sino Biological | 40592-V08H86 |
| SARS-CoV-2 Delta Spike | Sino Biological | 40589-V08B16 |
| SARS-CoV-2 Delta RBD | Sino Biological | 40592-V08H115 |
| SARS-CoV-2 Omicron BA.1 Spike | Sino Biological | 40589-V08H26 |
| SARS-CoV-2 Omicron BA.1 RBD | Sino Biological | 40592-V08H121 |
| SARS-CoV-2 Omicron BA.5 Spike | Sino Biological | 40592-V08H32 |
| SARS-CoV-2 Omicron BA.5 Spike | Sino Biological | 40592-V08H130 |
| SARS-CoV-2 Omicron BQ.1.1 Spike | Sino Biological | 40589-V08H41 |
| SARS-CoV-2 Omicron BQ.1.1 Spike | Sino Biological | 40592-V08H143 |
| SARS-CoV-2 XBB.1.5 Spike | Sino Biological | 40589-V08H45 |
| HCMV Glycoprotein B | Sino Biological | 10202-V08H1 |
| HCoV-OC43 Spike | Sino Biological | 40607-V08B1 |
| HCoV-NL63 Spike | Sino Biological | 40641-V07E |
| MERS-CoV Spike | Sino Biological | 40069-V08B |
| SARS-CoV-1 Spike | Sino Biological | 40634-V27H |
| Human FcγRIIA | Duke Human Vaccine Institute | Custom Order |
| Human FcγRIIB | Duke Human Vaccine Institute | Custom Order |
| Human FcγRIIIA | Duke Human Vaccine Institute | Custom Order |

| | | |
|---|---|---|
| Human FcγRIIIB | Duke Human Vaccine Institute | Custom Order |
| Human FcαR | Duke Human Vaccine Institute | Custom Order |
| Streptavidin-PE | Agilent Technologies | PB32-10 |
| Ebola Virus Glyoprotein | IBT Bioservices | 0501-015 |
| Influenza HA | Sino Biological | 11687-V08H |
| LC-LC-Sulfo-NHS Biotin | ThermoFisher | A35358 |
| Streptavidin-R-Phycoerythrin | Prozyme | PJ31S |
| **Software and algorithms** | | |
| GraphPad Prism 8 | GraphPad Software, Inc. | Ragon License |
| R Studio V. 4.0.4 | R Project for Statistical Computing | Open Source |
| Flow Jo | BD Bioscience | Ragon License |
| Python V 3.8.8 | MathWorks | Open Source |
| Matplotlib V 3.3.3 | Mathworks with Python | Open Source |
| **Other** | | |
| MagPlex microspheres | Luminex corporation | MC12001-01 |

**Supplementary Table 1. List of reagents and resources used in this study.**
